# Assessment and representation of variability in ecological niche model predictions

**DOI:** 10.1101/603100

**Authors:** Marlon E. Cobos, Luis Osorio-Olvera, A. Townsend Peterson

**Affiliations:** Department of Ecology and Evolutionary Biology & Biodiversity Institute, University of Kansas, Lawrence, Kansas 66045, USA; Facultad de Ciencias, Universidad Nacional Autónoma de México, Ciudad de México 04510, México

**Keywords:** hierarchical partitioning, model projection, model uncertainty, R function, species distribution models

## Abstract

Ecological niche models are popular tools used in fields such as ecology, biogeography, conservation biology, and epidemiology. These models are used commonly to produce representations of species’ potential distributions, which are then used to answer other research questions; for instance, where species richness is highest, where potential impacts of climate change can be anticipated, or where to expect spread of invasive species or disease vectors. Although these representations of potential distributions are variable which contributes to uncertainty in these predictions, model variability is neglected when presenting results of ecological niche model analyses. Here, we present examples of how to quantify and represent variability in models, particularly when models are transferred in space and time. To facilitate implementations of analyses of variability, we developed R functions and made them freely available. We demonstrate means of understanding how much variation exists and where this variation is manifested in geographic space. Representing model variability in geographic space gives a reference of the uncertainty in predictions, so analyzing this aspect of model outcomes must be a priority when policy is to be set or decisions taken based on these models. Our open access tools also facilitate post modeling process that otherwise could take days of manual work.

## INTRODUCTION

Ecological niche modeling (ENM) has become a tool set with extremely high interest and attention in the past two decades (Peterson et al. 2011). The main aim of these models is to characterize environmental conditions that allow a species to maintain populations at a site. By characterizing suitable conditions, these methods allow researchers to predict, among other things, where a species can live on the planet, or what would happen to that species if environmental conditions change (Evans 2012). As such, these methods have become popular in studies attempting to understand species’ current distributions, processes of invasion, disease risk explorations, or predict impacts from climate change on species’ distributions (Peterson 2006; Yates et al. 2018). Although such analyses offer tempting predictions, they are based on occurrence data and climate models, which has drawn criticism owing to the inherent—and largely unassessed—uncertainty that they manifest (Dobrowski et al. 2011).

Uncertainty in ENMs can derive from multiple sources (Heikkinen et al. 2012; Petchey et al. 2015). Recent theoretical and methodological advances have allowed development of novel approaches for identifying adequate complexity levels during model calibration (i.e., testing distinct sets of predictors, response types, and fitting/overfitting; Warren et al. 2010; Muscarella et al. 2014). Since more than one parametrization may result in adequate models after calibration (Gupta et al. 1998), using multiple parameter settings seems logical (Cobos et al. 2019). This consideration of multiple sets of parameters adds a further important source of variation in model outcomes (Peterson et al. 2018). One current common application of ENMs is projecting them in time to understand potential future changes in species’ distributions (Qiao et al. 2018). This practice usually implies the use of general circulation model (GCM) outputs and emissions scenarios (representative concentration pathways, or RCPs), adding still more sources of variation (Diniz-Filho et al. 2009). As a consequence, uncertainty in model projections is higher in these transferability applications (Elith and Leathwick 2009; Yates et al. 2018).

Despite the existence of variability in ENMs, assessing variability and presenting visualizations of uncertainty is not a frequent practice (Peterson et al. 2018, but see Dörmann et al. 2008; Lemes and Loyola 2013). This gap in the field is potentially a consequence of the difficulty of measuring and showing variability when various sources of variation are considered and when they are manifested on multiple levels (Yates et al. 2018). Indeed, assessment and representation of model variability may take days when performed manually. As a consequence, no consensus exists on how to assess and represent variability in ENMs, and examples of uncertainty analysis in the literature are not common (Beale and Lennon 2012).

Hence, here, we aimed to (1) illustrate how to analyze and represent model variability produced from multiple sources of variation in ENMs; and (2) implement open source tools (R functions) that allow performing such analyses and representations. As case studies, we use two species with distinct distributional characteristics: a tick broadly distributed in North America, *Amblyomma americanum* (L., 1758), and a toad endemic to the Cuban archipelago, *Peltophryne empusa* Cope, 1862.

## METHODS

### Example species

Species selected as examples were selected owing to their different ecological and distributional features which lead to contrasting results regarding model variability, one species restricted to an archipelago and the other more broadly distributed in the continent. Occurrence and environmental data for the two example species used in this study have been previously used in the presentation of the kuenm R package (Cobos et al. 2019), and are openly available at https://doi.org/10.17161/1808.26376. These data set is as follows: for the tick, 178 filtered occurrences, divided in 89 records for training and 89 for testing; for the toad 64 filtered records, where 50% of the occurrences were used for training and 50% for testing. These data were originally derived from 14,831 records for the tick, and from 230 for the toad, but have been thinned and reduced to avoid negative effects of spatial autocorrelation (for details on thinning process, see Raghavan et al. 2019; Cobos et al. 2019). Environmental data sets used in our example comprise 4 “bioclimatic” variables for each species (spatial resolution = 10’ for the tick, and 30” for the toad) and were drawn from the WorldClim database v1.4 (Hijmans et al. 2005; available at http://www.worldclim.org/current). Variables used for each analysis were annual mean temperature, annual precipitation, precipitation seasonality, and precipitation of driest quarter, for the tick; and temperature seasonality, minimum temperature of coldest month, temperature annual range, and precipitation of wettest month, for the toad. Variables were selected via rigorous processes of model calibration in which distinct parameter settings as well as alternative sets of variables were tested (see Raghavan et al. 2019; Cobos et al. 2019).

### Ecological niche modeling

Models were generated with 10 replicate resamplings (bootstrap), using Maxent (Phillips et al. 2006) and the results of model calibration obtained from application of the kuenm R package (Cobos et al. 2019). Model calibration consisted in evaluating models created with 19 distinct regularization multipliers (0.1 to 1 at intervals of 0.1, 1 to 6 at intervals of 1, and 8 and 10), 29 feature classes (resulted from all combinations of linear, quadratic, product, threshold, and hinge response types), and three distinct sets of environmental variables with distinct number of variables each. Best parameter settings were selected considering statistical significance (partial ROC; Peterson et al. 2008), predictive power (omission rates *E* = 5%; Anderson et al. 2003), and complexity level (AICc; Warren et al. 2010), in that order (Table 1; Cobos et al. 2019). For the tick, models were calibrated in across the United States and projected to all the world in current and future scenarios. We calibrated the toad models across the whole Cuban archipelago, and projected them to future scenarios for this country. Model projections were performed allowing extrapolation with clamping in predictions, although these assumptions (i.e., to allow or not extrapolation) can be changed in other applications. Future conditions were characterized by three climate models (GCMs; BCC-CSM1-1, CCSM4, and MIROC5) and two greenhouse gas emissions scenarios (RCP; 4.5 and 8.5; IPCC 2013). Future environmental datasets were not part of the data used in Cobos et al. (2019); they were obtained from the WorldClim database v1.4 (available at http://www.worldclim.org/CMIP5v1).

**Table 1.**
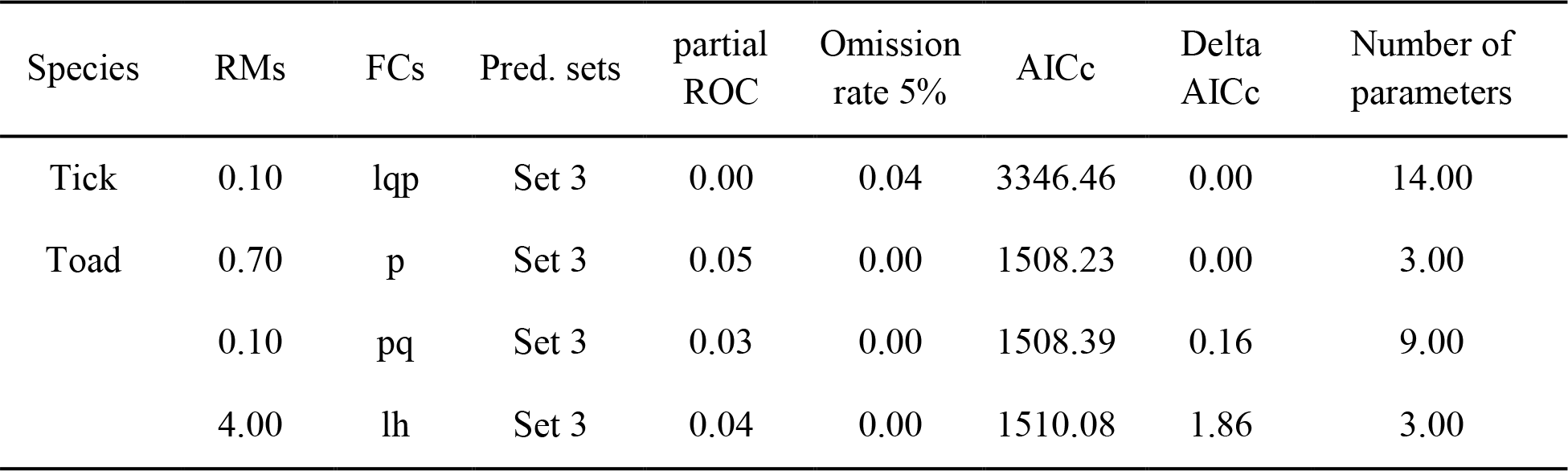
Performance metrics for parameter settings regarding regularization multiplier (RMs), feature classes (FCs), and sets of environmental predictors (Pred. sets), used for creating final models for the two example species. FCs are as follows: linear = l, quadratic = q, product = p, and hinge = h. Modified from Cobos et al. (2019).

| Species | RMs | FCs | Pred. sets | partial<br>ROC | Omission<br>rate 5% | AICc | Delta<br>AICc | Number of<br>parameters |
| --- | --- | --- | --- | --- | --- | --- | --- | --- |
| Tick | 0.10 | lqp | Set 3 | 0.00 | 0.04 | 3346.46 | 0.00 | 14.00 |
| Toad | 0.70 | p | Set 3 | 0.05 | 0.00 | 1508.23 | 0.00 | 3.00 |
|  | 0.10 | pq | Set 3 | 0.03 | 0.00 | 1508.39 | 0.16 | 9.00 |
|  | 4.00 | lh | Set 3 | 0.04 | 0.00 | 1510.08 | 1.86 | 3.00 |

Model projections for the tick therefore had three sources of variation (replicates, GCMs, RCPs) and model projections for the toad had the same three sources plus parameter settings because more than one set of parameters was selected as optimum for this species. We did not evaluate final models to maximize sample sizes available for uncertainty analysis, although such analyses should be inherent in all modeling efforts designed for decision making. All processes of model development were performed using the kuenm R package, which uses Maxent as the modeling algorithm.

### Assessment and representation model variability

Four main post-modeling analyses/processes were used to assess and represent model variability in the two examples: (1) calculation of model statistics (i.e., median and range), (2) identification of changes of suitable areas and suitability in future projections, (3) creation of maps showing the variance contributed by each source of variation, and (4) hierarchical partitioning of the variance in models among sources of variation. The four sources of variation considered in these analyses were: model replicates, parameter settings (parameters), GCMs, and RCPs, although one could also use different algorithms, which would add another source of variation (Qiao et al. 2015). We followed routines similar to those proposed by Campbell et al. (2015) for creating maps of changes of suitable areas, and by Peterson et al. (2018) for mapping variance and exploring results of the hierarchical partitioning of the variance. All of our analyses were performed in R, version 3.5.1 (R Core Team 2018).

Descriptive statistics for ENMs are usually calculated by Maxent as part of models created with replicates. However, sometimes, more than one parameter setting needs to be considered when developing these models (Muscarella et al. 2014; Radosavljevic and Anderson 2014; Cobos et al. 2019); as such, new calculations need to be developed to represent model outcomes. These new calculations are needed for creating consensus among results from distinct parameterizations, but also for showing how variable results are depending on the parameters used. Such was the case for the toad species: we calculated the median and range of all model replicates across all parameter settings, distinguishing among distinct areas of projection (i.e., calibration area and area of future projection).

Changes in future projections were identified using model statistics calculated in the previous step and considering patterns of agreement among GCMs when detecting zones of stability, gain, and loss of suitable areas. All analyses were done separating future projections by the two RCPs considered. To identify changes in suitability, we calculated the difference between the median of all future models constructed with distinct GCMs and the model for the current period: positive values indicate increases in suitability and negative values represent decreases in suitability. Analyses to identify changes in suitable areas consisted of the following steps: (1) threshold all future median models (0 = unsuitable, 1 = suitable); (2) threshold the current median model to values of 0 = unsuitable versus 1 + total number of GCMs = suitable; (3) sum the current binary with all future binary models; and (4) identify areas of stability, gain, and loss of suitable areas. This last step was interpreted as follows: areas with values of 0 represent stable unsuitable areas in all GCMs; values from 1 to the total number of GCMs indicate gain of suitable areas in same number of GCMs; values from {1 + [(total number of GCMs × 2) − 1]} to (1 + total number of GCMs) indicate loss of suitable areas in one to all the GCMs, respectively; and values equal to (total number of GCMs × 2) indicate stable suitable areas in all GCMs.

Maps of variance were created for each source of variation considered in models. The process consisted of calculating the mean of all the models representing each level in each source of variation and then calculating the variance across these means. For instance, in a model projected to future conditions, all first replicates of models created with distinct parameter settings, GCMs, and RCPs, were put together; we calculated the mean of this set of layers and repeated the process for all other replicates. After that, we calculated the variance of the 10 layers representing the means of the 10 replicates. The resulting layer was used to represent model variability coming from replicates; similar routines were performed for other sources of variation.

We estimated statistically the amount of variance coming from distinct sources of variation in our model outcomes using the hierarchical partitioning analysis (Chevan and Sutherland 1991) implemented in the R package hier.part (Walsh and Mac Nally 2013). Specially, we extracted the values from each model using a random sample (*N* = 1000) and arranged those data for identifying all sources of variation involved (i.e., replicates, parameters, GCMs, RCPs). The resulting matrix was used to perform the hierarchical partitioning analysis. To detect whether total effects detected for each source of variation were statistically different than zero, we repeated this process 100 times, using distinct random samples, and calculated the 95% confidence intervals (CIs). Statistically significant results were those that did not include zero in their intervals.

### Open access tools

We implemented all the above routines as R functions. These functions were constructed to help users in handling the sometimes-overwhelming number of results from modeling exercises. These functions were written to be useful in diverse situations and to avoid potential problems related to RAM usage in personal computers with low computational and memory capacity. Although these examples were created with only one option of extrapolation settings (i.e., allowing extrapolation with clamping), functions are developed to be able to handle other settings regarding extrapolation, or all of them in a loop. R packages used in these functions are raster (Hijmans et al. 2017), rgdal (Bivand et al. 2018), and hier.part.

## RESULTS

Models for the two example species represented their potential distributions adequately as they are not overfit to geographic records and omission is only related to the threshold selected. For the tick, in the calibration area, the range and variance maps had the same geographic pattern, though their values were different (Figure 1). In the worldwide projection for present, suitable areas occupied tropical and subtropical areas. Range and variance maps from replicates had similar patterns as in the calibration area (Figure 1).

**Figure 1.**
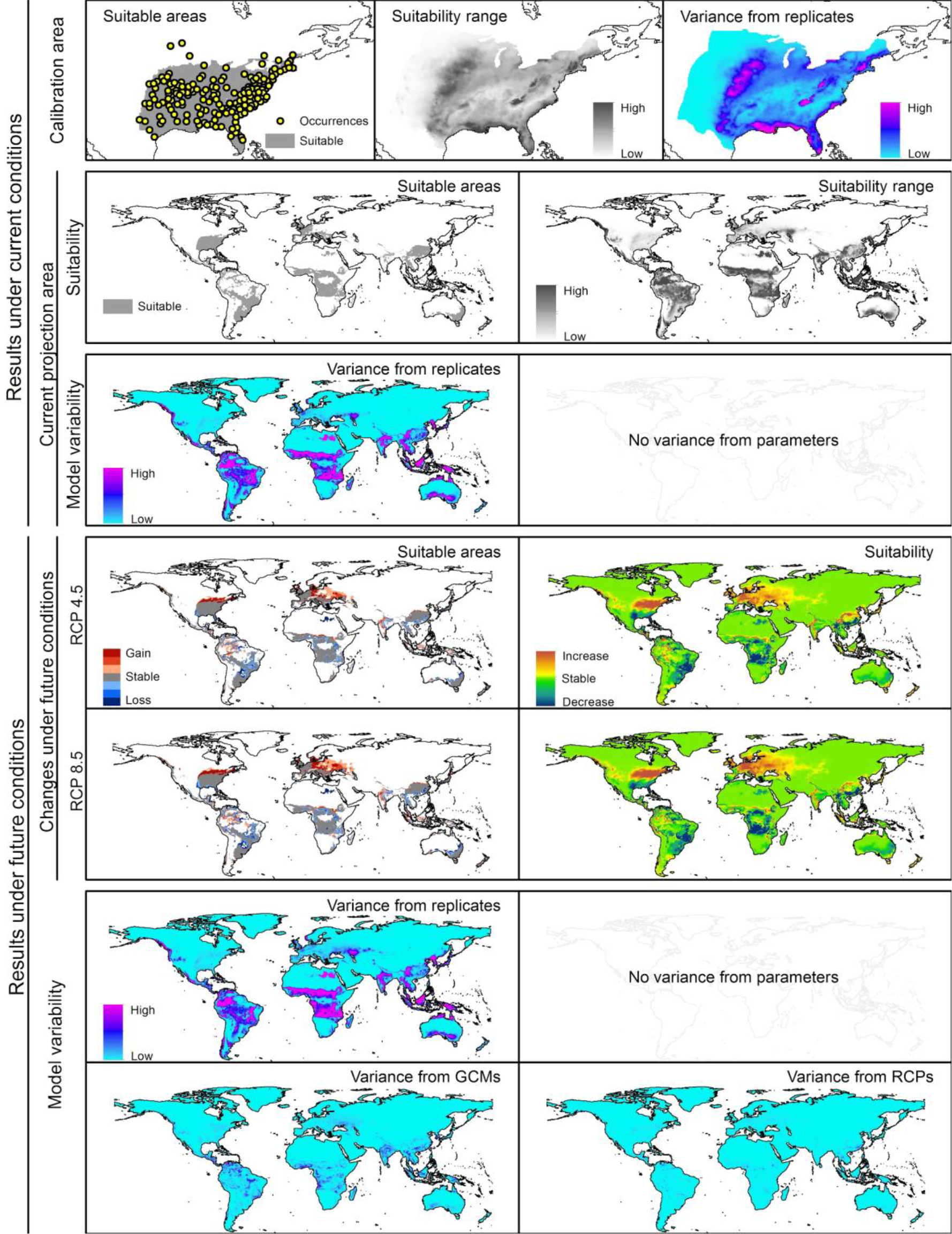
Summary of ecological niche modeling and variation assessment results for the tick species.

As regards of changes in suitable areas and suitability values in future, for the tick, patterns of changes were similar between the two RCPs. For RCP 8.5, gained suitable areas were slightly bigger in northern parts of Europe and North America; more changes in suitability values were observed in the RCP 8.5 projections than in the RCP 4.5 projections (Figure 1). Variance maps indicated that model replicates were the main source of variation in future projections, and higher variability were found in tropical areas for all sources of variation, and in some Eurasian areas for replicates (Figure 1). Patterns detected with hierarchical partitioning agreed with those found in variance maps. Effects of RCPs on overall variance were the smallest of the four factors, although their contribution was statistically different from zero (Figure 3).

For the toad, in the current period (note that because this model was calibrated across the entire Cuban archipelago, no current projection was needed), suitable areas occupied almost all lowland areas in the country (Figure 2). Maps of model range and variance from replicates had similar patterns, as in the previous species. In this case, however, another source of variation existed, parameters, as multiple optimal models were detected; parameters turned out to show the highest levels of variance. Variance from parameters did not have patterns similar to those found for replicates or suitability range (Figure 2). The hierarchical partitioning analyses confirmed these patterns: parameters constituted around 95% of total effects on the variance. Effects from replicates, although smaller, were still statistically significant (Figure 3).

**Figure 2.**
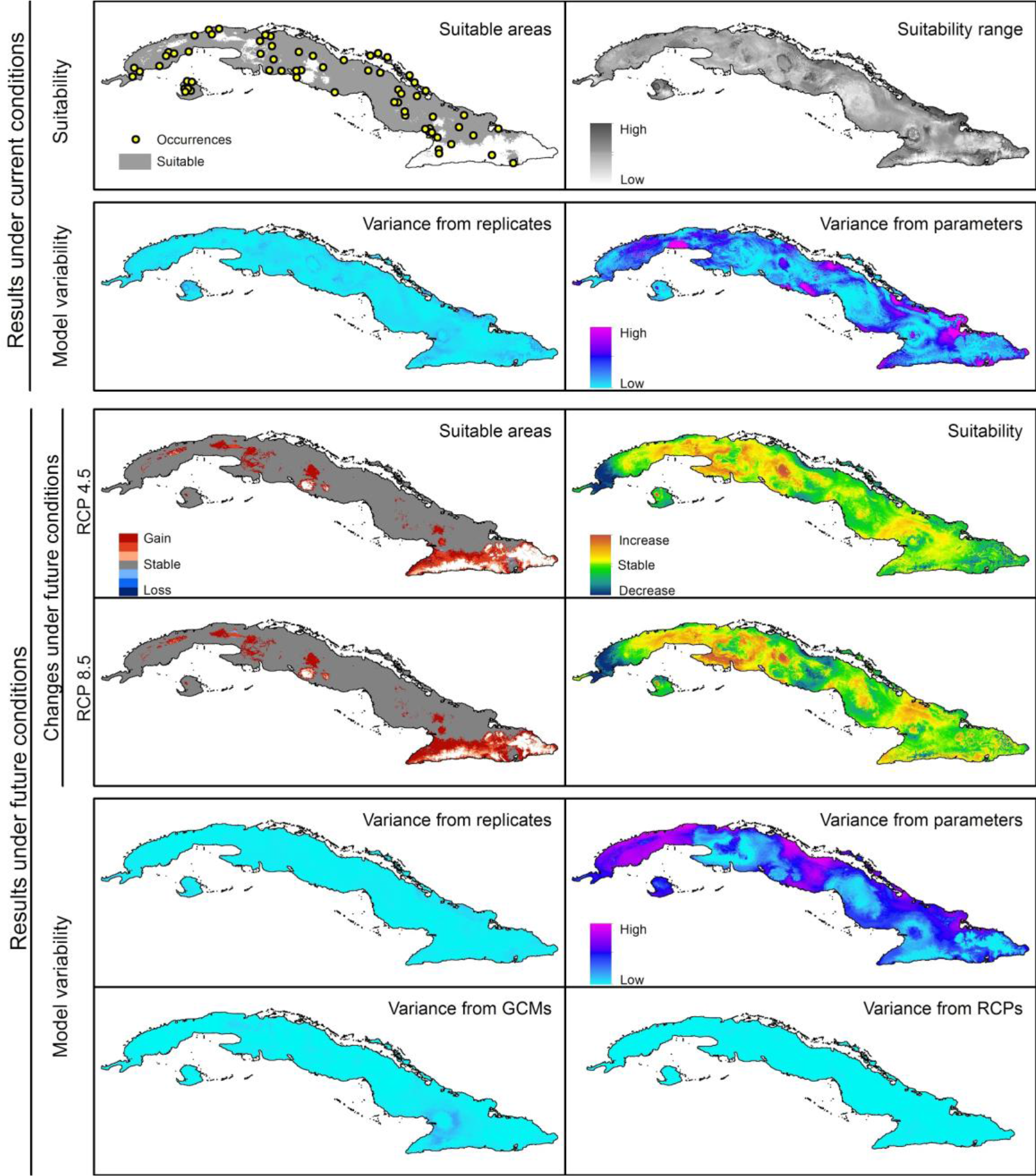
Summary of ecological niche modeling and variation assessment results for the toad species.

**Figure 3.**
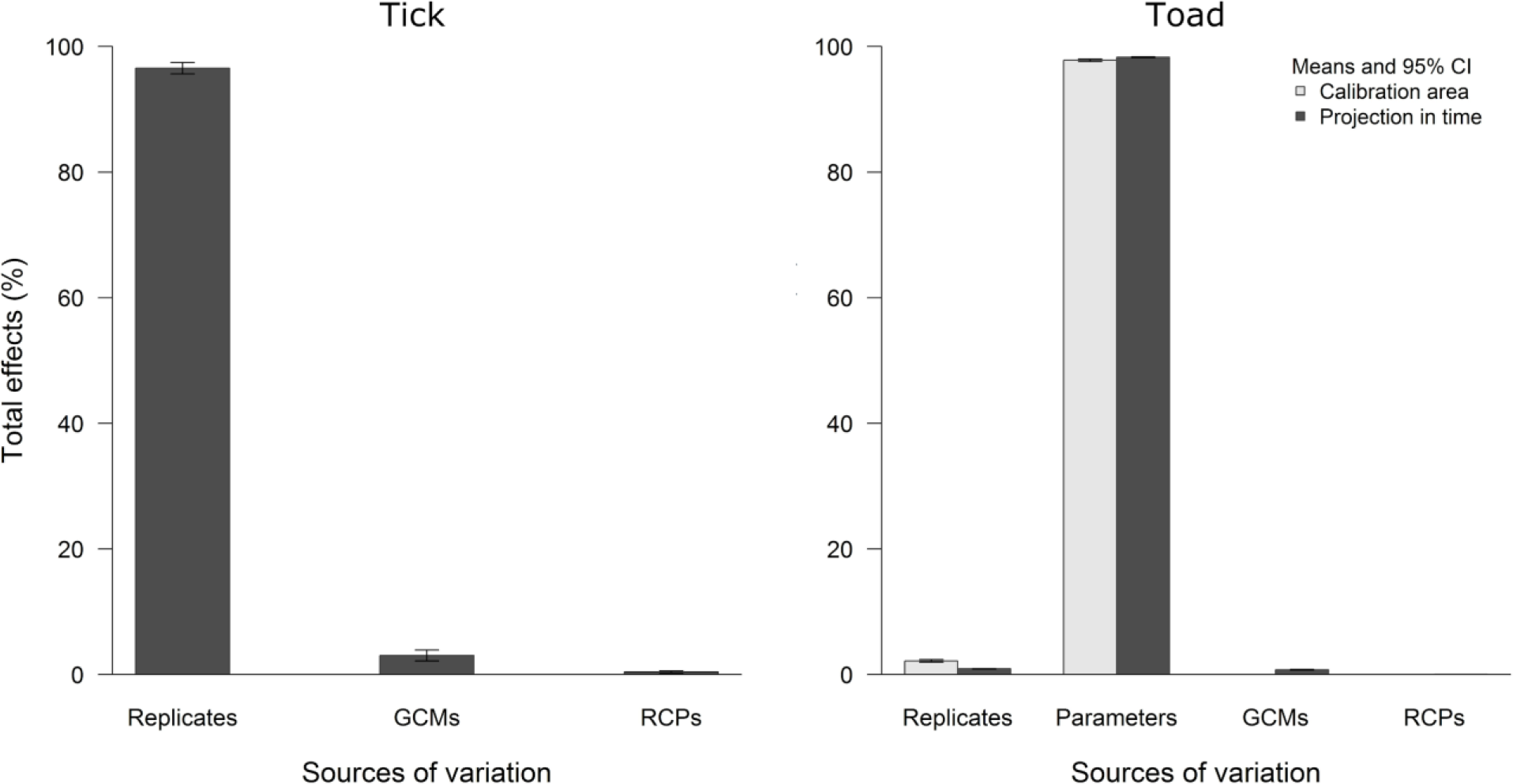
Results of the hierarchical partitioning of variance deriving from distinct sources of variation in ecological niche models for the two example species (CI = confidence intervals).

The two RCPs revealed closely similar patterns regarding changes in future suitability of areas for the toad. Most changes detected in future suitable areas were gains, especially in areas of medium-to-high elevation, with slightly larger areas under RCP 8.5 than under RCP 4.5. Suitability values, as for the previous species, changed slightly more in RCP 8.5; geographic patterns of change in these values were similar but not totally coincident (Figure 2). Maps of variance for future projections of models indicated that parameters were the source that contributed the most to overall model variability. Highest values of variance were found in the western and central parts of the archipelago, with a fragmented pattern. Other sources contributed little to variability in future projections of models. In fact, the only one with a detectable pattern was GCMs, with the highest variance values in the eastern part of Cuba (Figure 2). The hierarchical partitioning results allowed us to see the even-higher effects of distinct parameter settings on variance in future projections than in the current period. Replicates and GCMs had small effects, but their CIs at 95% did not include zero; the effect of RCPs, however, was not distinguishable from zero (Figure 3).

The set of tools created to carry out the analyses performed well, even when computationally challenging tasks were executed all, without excessive demands on the computer’s RAM. The use of these functions helped considerably in handling the large number of model outputs considered in the analyses (180 for the tick, 360 for the toad). All main and helper functions developed for performing analyses (Table 2) are implemented in the new version of the kuenm R package (Cobos et al. 2019, available at https://github.com/marlonecobos/kuenm).

**Table 2.** Description of the main R functions created for assessing and presenting variation in ecological niche models.

| Function name | Description |
| --- | --- |
| kuenm_modstats | Model descriptive statistics calculation (raster inputs and outputs). Calculates metrics of position and dispersion (i.e., median, mean, variance, standard deviation, minimum, maximum, and range) of model outputs across multiple parameter settings. |
| kuenm_projchanges | Calculation of agreement of model changes predicted using multiple GCMs in future projections (raster inputs and outputs). |
| kuenm_modvar | Identification of geographic patterns of variability in model predictions (raster inputs and outputs). |
| kuenm_hierpart | Statistical assessment (i.e., hierarchical partitioning) of variation in model predictions (raster inputs, graphical and numerical outputs). |

## DISCUSSION

### Example outcomes

In general, exploring our examples allowed us to visualize how model variability can give an idea of the uncertainty in our predictions. Areas where variability was highest and GCM agreement was low (Figures 1 and 2) were more difficult to interpret, and any potential conclusions about such areas should be made with caution. The two examples had in common a low contribution to the overall variance by GCMs and RCPs (Figure 3). Considering what these two sources represent, if we think only about GCMs, the outcome is good because that would mean that using distinct climate models will not change the general pattern of future changes (but see Diniz-Filho et al. 2009 p.; Tang et al. 2018). However, if we think about what RCPs are representing (distinct global scenarios of greenhouse gases; IPCC 2013), we could have expected more variation coming from this source. Replicates contributed importantly to the variance in the tick models, but not for the toad results. Parameters, however, had the biggest effects on model variability, which means that careful attention must be paid to model selection and exploration of large numbers of candidate models.

We know of few examples in the literature in which model variability has been assessed using variance partitioning approaches (e.g., Ramírez-Gil et al. in review; Dörmann et al. 2008; Lemes and Loyola 2013; Peterson et al. 2018). In those cases, GCMs were important sources of variation; replicates, on the contrary, were not as important regarding variance contribution. RCP contributions were lower than those of GCMs. Only two of these previous studies analyzed parameters as sources of variation; in both of them, this source had the largest effects on model variability.

### How representing model variability is useful?

Representing model variability and agreement in predictions of potential future change is challenging (Yates et al. 2018). This complexity derives mainly from the time that the post-modeling processing demands, especially when working with large numbers of model outputs that result from combinations of distinct replicates, parameter settings, climate models, and emission scenarios, when creating models and their projections. Presenting model variability, however, allows detecting important characteristics of model outcomes that would otherwise be neglected (i.e., how variable are models and where is this variability higher; Gould et al. 2014).

Agreement of different predictions potential change in distributions in the future, for example, may play a key role in selecting priority areas for conservation (e.g., areas where potential refugia for various taxa are identified in the face of climate change with greater confidence; Morelli et al. 2016; Robinson and Fordyce 2017). On the other hand, detecting areas where high levels of variability are present may be useful for establishing programs for monitoring changes given the high uncertainty that they present. Potential uses of the information on model variability—directly related to uncertainty in model predictions—is therefore a key element of what researchers and stakeholders should consider before making conservation-related decisions (Gregr and Chan 2015; Sequeira et al. 2018).

Assessing how much distinct sources contribute to overall variability, as well as the geographic pattern of this variability, also helps to understand how distinct factors may affect model predictions. Model variability for distinct species would receive higher or lower contributions from replicates, parameters, or other sources, depending on diverse factors. That is, replicates may contribute more than parameters for some species, but the opposite may be the case for others. We have not detected a clear relationship between the number of classes of the source of variation and the amount of variation contributed by it. In fact, for one of the two example species analyzed here, parameters (*N* = 3) contributed more than replicates (*N* = 10). We therefore suggest that variance contributions likely depend on multiple factors; for instance, results may change depending on the environmental heterogeneity of the calibration area, or on how numerous are the species records and how are they distributed in the environmental space. Each species, therefore, constitutes an individual and unique case, and assumptions should not be made without performing proper analyses.

### Implications of using the new tools

The set of tools developed for this study (i.e., R functions) offer convenient options for performing post-modeling processes to assess and represent model variability. The main challenge for researchers when trying to use these tools is organizing their data before starting (for a detailed explanation see https://github.com/marlonecobos/kuenm_model_var_functions). However, this step is unnecessary if modeling processes are performed using the kuenm R package, as kuenm outputs are formatted automatically for the present set of functions. Although performing all these analyses still can take several hours in a computer with common features, the time for organizing the data (if required) and preparing the few lines of necessary code (less than an hour) is not comparable with the days of manual processing previously needed. Perhaps the most advantageous aspect of using these tools is that processes can be automatized for performing analyses for multiple species.

### Concluding remarks

A better representation of model variability would, without doubt, improve the way in which we interpret ecological niche models, especially in geographic spaces. However, the new implementations presented here do not replace the use of other analyses used to measure uncertainty when creating ENMs (e.g., analyses of uncertainty derived from quality of data; Gould et al. 2014). To the contrary, the analyses presented here complement the other metrics such that researchers can explore limits of interpretation for conclusions from ENMs. Our results showed that variability in models could come in distinct amounts from various sources, and that geographic patterns of model variability are not easily deducible. An important consideration to add is that distinct types of extrapolation will also influence results regarding variance considerably (Owens et al. 2013). That is, allowing free extrapolation in model projections could change the proportion of variance contributed by replicates or GCMs, compared to those contributions in model projections in which extrapolation was not allowed.

Tools for measuring variability in model predictions are scarce (but see Gould et al. 2014); our set of tools makes measuring variability related to model disagreement more straightforward. The functions presented here have been designed to work with four sources of variation but do not include distinct algorithms. Our reasoning for not including this source of variation is that distinct algorithms generate different outcomes, and these outcomes are not necessarily comparable; however, given the potential for using consensus models procedures (Zhu and Peterson 2017), further such implementations could take this dimension into account in the future

## ACKNOWLEDGMENTS

The University of Kansas supported MEC during the period in which this project was developed. R. Ravaghan allowed us using his data for one of the examples of application. A. Alkishe helped us with testing our R functions and gave us inputs on how to improve them.

## Author Contribution

MEC and ATP designed the analyses and tools. MEC and LOO implemented and programmed the R tools. MEC wrote the manuscript. All authors reviewed the manuscript and gave final approval for publication.

